# Molecular Survey for Selected Viral Pathogens in Wild Leopard Cats (*Prionailurus bengalensis*) in Taiwan with an Emphasis on the Spatial and Temporal Dynamics of Carnivore Protoparvovirus 1

**DOI:** 10.1101/2020.02.21.960492

**Authors:** Chen-Chih Chen, Ai-Mei Chang, Wan-Jhen Chen, Po-Jen Chang, Yu-Ching Lai, Hsu-Hsun Lee

## Abstract

The leopard cat (*Prionailurus bengalensis*) has been listed as an endangered species under the Wildlife Conservation Act in Taiwan since 2009. In this study, we targeted viral pathogens, included carnivore protoparvovirus 1 (CPPV-1), feline leukemia virus (FeLV), feline immunodeficiency virus (FIV), coronavirus (CoV), and canine morbillivirus (CMV), using molecular screening. The spatial and temporal dynamics of the target pathogens were evaluated. Through sequencing and phylogenetic analysis, we aimed to clarify the phylogenetic relationship of isolated viral pathogens between leopard cats and domestic carnivores. Samples from 23 and 29 leopard cats that were live-trapped and found dead, respectively, were collected from Miaoli County from 2015 to 2019 in northwestern Taiwan. CPPV-1 and coronavirus were detected in leopard cats. The prevalence (95% confidence interval) of CPPV-1, and CoV was 63.5% (50.4%–76.6%) and 8.8% (0%–18.4%), respectively. The majority of sequences of each CPPV-1 strain amplified from Taiwanese leopard cats and domestic carnivores were identical. All the amplified CoV sequences from leopard cats were identified as feline coronavirus. The spatial and temporal aggregation of CPPV-1 infection in leopard cats was not determined in the sampling area, which indicated a wide distribution of CPPV-1 in the leopard cat habitat. We consider sympatric domestic carnivores to be the probable primary reservoir for the pathogens identified. We strongly recommend establishing efforts to manage CPPV-1 and FCoV in the leopard cat habitat, with an emphasis on vaccination programs and population control measures for free-roaming dogs and cats.

**IMPORTANCE:** The leopard cat (*Prionailurus bengalensis*) is an endangered species in Taiwan. The effects of infectious diseases on the wildlife population have increasingly been recognized. In this study, we targeted highly pathogenic viral pathogens in wild cat species, included carnivore protoparvovirus 1 (CPPV-1), feline leukemia virus (FeLV), feline immunodeficiency virus (FIV), coronavirus (CoV), and canine morbillivirus (CMV), using molecular screening. Furthermore, we collected the epidemiological and phylogenetic data to understand the spatial and temporal dynamics of the target pathogens in the wild leopard cat population and identified the possible origin of target pathogens. Based on our study, we consider sympatric domestic carnivores to be the probable primary reservoir for the pathogens identified. Our study provides a deeper understanding related to the distribution of target viral pathogens in the wild leopard cats. The information is essential for leopard cat conservation and pathogen management.

## INTRODUCTION

The leopard cat (*Prionailurus bengalensis*) is an endangered felid species that is distributed in East, Southeast, and South Asia (1). It was previously commonly distributed in the lowland habitats throughout the island of Taiwan (2, 3). However, the Wildlife Conservation Act of Taiwan listed the leopard cat as an endangered species in 2009 after an island-wide decline in the population of this species (4). Currently, the distribution of Taiwanese leopard cats is restricted to small areas in 3 counties in Central Taiwan, namely Miaoli, Nantou, and Taichung City. Studies in Miaoli County suggested that road traffic, habitat fragmentation and degradation, illegal trapping, and poisoning are the principal threats to the sustainability of the leopard cat population (5). However, the possible direct or indirect effects of pathogens on the population of Taiwanese leopard cats have never been evaluated. Moreover, information related to infectious agents distributed in the wild Taiwanese leopard cat population has remained scarce. Our previous study documented the distribution of carnivore protoparvovirus 1 in Taiwanese leopard cats and its association with domestic carnivores (6). To our knowledge, this was the only study on infectious agents in free-living leopard cats in Taiwan. The effects of infectious diseases on the wildlife population have increasingly been recognized (7, 8). Conspicuous illness or the mass die-off of wild animals caused by specific agents are easier to identify and are usually considered a threat to the abundance of wildlife populations. Although unremarkable or sublethal diseases in wild animals are difficult to identify, such diseases may reduce the fitness of wild animals through an increased energy output or decreased food ingestion, arresting the growth of the population substantially (7, 9).

Pathogen infection in wild felids has been documented worldwide with different degrees of importance. Viral pathogens that have been identified in wild or captive leopard cats include feline immunodeficiency virus (FIV) (10), carnivore protoparvovirus 1 (CPPV-1) (6, 11, 12), feline herpesvirus type 1 (FHV-1) (11), and feline calicivirus (FCV) (11). Furthermore, studies have recorded infection by bacterial and parasitic agents including *Anaplasma* (13, 14), hemoplasma (13, 15), *Hepatozoon felis* (16–18), and several helminths (19). Although the effects of the recorded infectious agents on leopard cats remain unclear, identifying infectious agents in the leopard cat population is essential for disease management and species conservation.

Our previous study recorded carnivore protoparvovirus 1 (CPPV-1) infection in free-living leopard cats, albeit with a limited sample size. In the present study, we extended the target of viral pathogens for screening using a larger sample size. The target viral pathogens were CPPV-1, feline leukemia virus (FeLV), FIV, coronavirus (CoV), and canine morbillivirus (CMV).

Our objective was to identify the infection of selected viral pathogens based on molecular screening. The spatial and temporal distribution of target pathogens was described. Furthermore, through sequencing and phylogenetic analysis, we aimed to clarify the phylogenetic relationship of isolated viral pathogens between leopard cats and domestic carnivores.

## MATERIALS AND METHODS

### Study area

All the leopard cats samples were collected from Miaoli County in northwestern Taiwan (Fig. 1). The sampling area has a hilly landscape with an elevation of less than 320 m above sea level. The total area of Miaoli County is 1820 km^2^, consisting of 1245.3 km^2^ of forests (68.8%), 291.2 km^2^ of agricultural land (16.1%), and 132.6 km^2^ of human construction (7.3%). A well-developed road system, which includes a primary road (approximately 25 m wide), secondary roads (approximately 10 m wide), and tertiary roads (approximately 5 m wide), and human encroachment have fragmented the wildlife habitat in this rural area. The Taiwanese leopard cat population was primarily distributed in the west half of Miaoli County (20).

**FIG 1.**
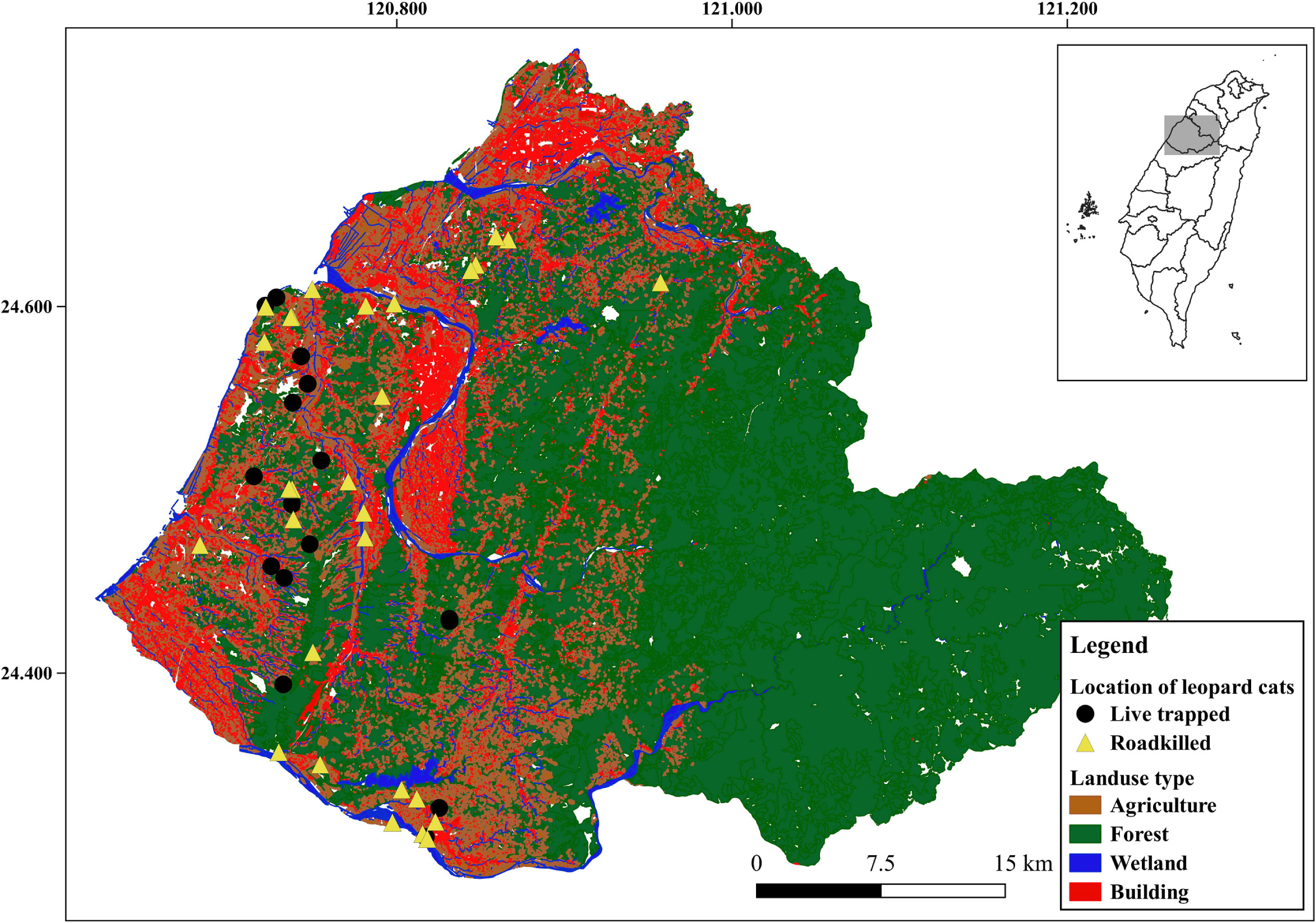
Sampling sites of leopard cats in Miaoli County. The map of Taiwan in the box indicates the location of Miaoli County in Taiwan. Circles and triangles, respectively, denote leopard cats that were live-trapped and found dead. Distribution of land-use types, comprising agriculture, forest, wetland, and building area, are denoted in the background.

Although estimates of the population of stray or free-roaming dogs and cats were not available, they were commonly observed and were sympatric with the leopard cats in the study area (20).

### Sample collection

The leopard cat samples were collected from January 2015 to April 2019. Free-living leopard cats were trapped for radio telemetry tracking or relocation of leopard cats that invaded poultry farms. Permission for conducting this study was issued by the Forest Bureau (Permit no.: COA, Forestry Bureau, 1061702029, 1081603388). Steel-mesh box traps (108-Rigid Trap, Tomahawk Live Trap, LLC., Hazelhurst, Wisconsin, USA) baited with live quails were employed for trapping the leopard cats. The trapped leopard cats were anesthetized by veterinarians using a mixture of dexmedetomidine hydrochloride (100 µg/kg) and tiletamine HCl/zolazepam HCl (2 mg/kg). The procedures for leopard cat trapping, anesthesia administration, and sample collection were approved by the Institutional Animal Care and Use Committee of National Pingtung University of Science and Technology (Approval no.: NPUST-106-014, NPUST-107-041).

The carcasses of found-dead (FD) leopard cats, with the majority of deaths caused by vehicle collision, were collected and submitted by the County Government of Miaoli for additional necropsy and sample collection.

Tissues and swabs collected for PCR or reverse transcriptase (RT-PCR) screening of selected pathogens are displayed in Table 1.

**TABLE 1.**
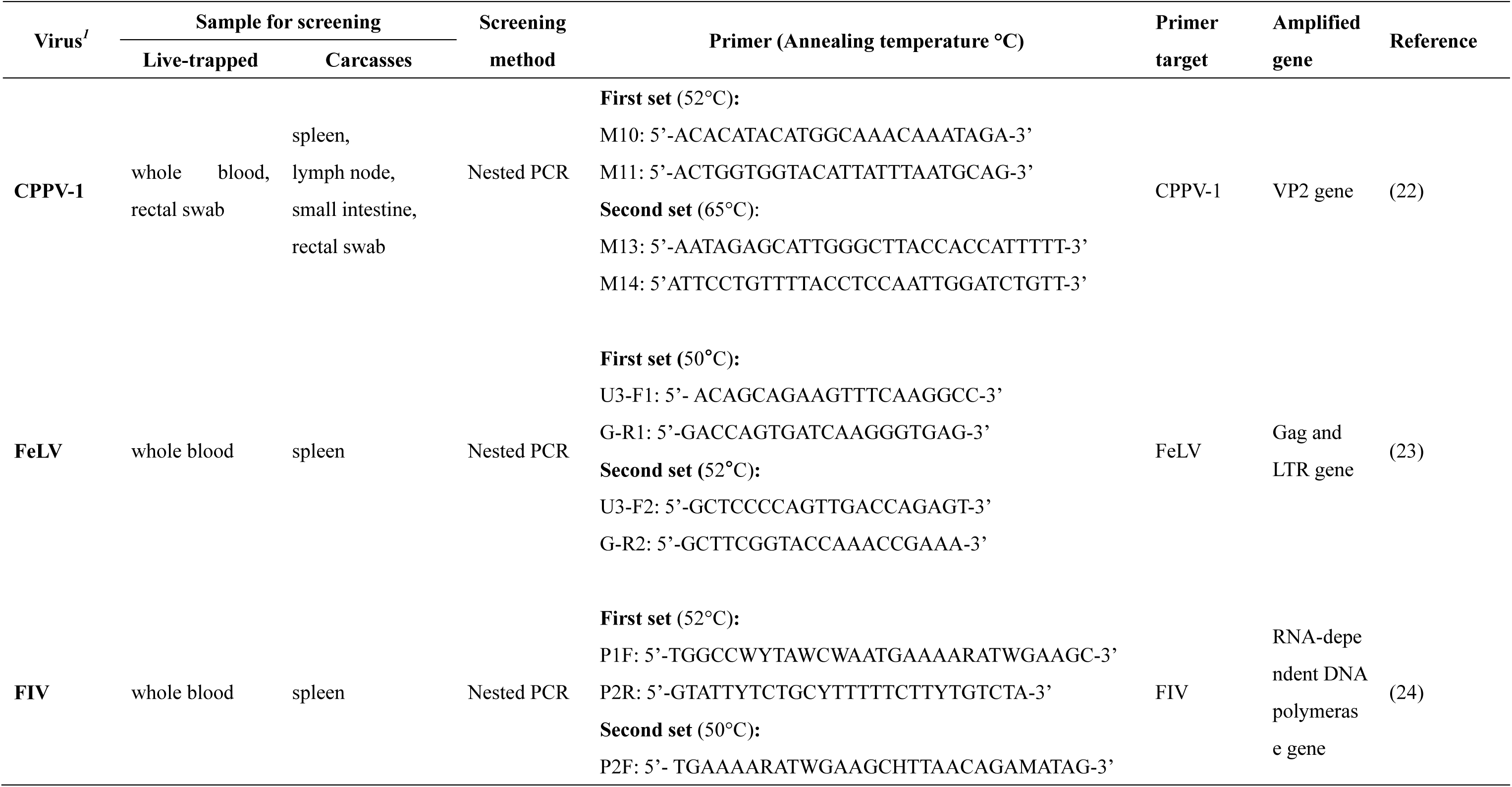
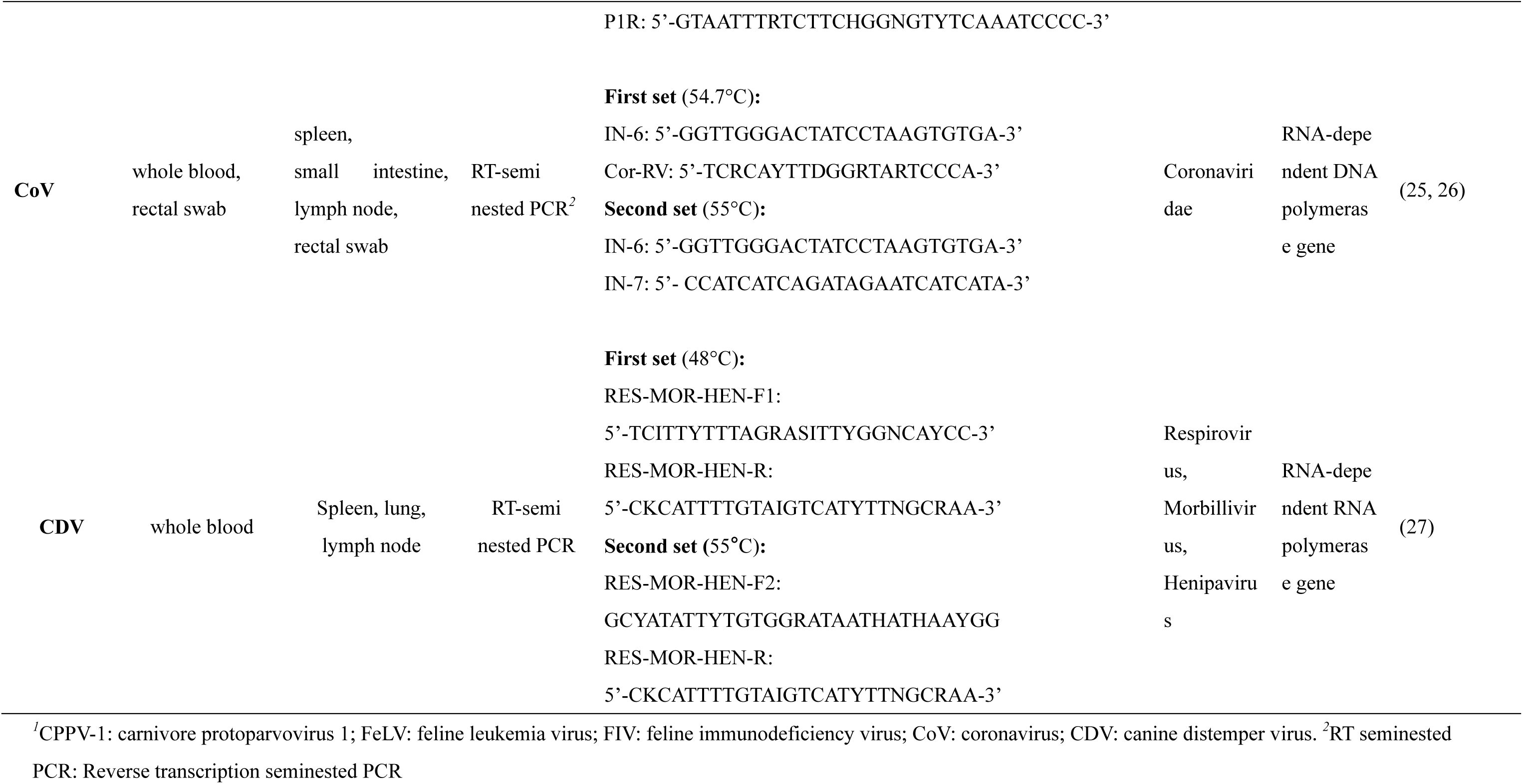
Samples collected from free-living leopard cats and PCR primers used for amplifying the target pathogens

We recorded sex and age for each leopard cat. Age classification was based on guidelines from Chen et al. (6). The criteria of age classification were deciduous dentition for juveniles, permanent dentition but not full growth for subadults, full growth of permanent dentition to mild abrasion of canine teeth for young adults, and moderate to severe abrasion of canine teeth for old adults.

### Nucleic acid extraction and (RT)PCR screening for selected viral pathogens

Samples were homogenized prior to nucleic acid extraction. Total DNA was extracted from the collected tissues and blood samples using the DNeasy blood and tissue kit and total RNA was extracted using the RNeasy minikit and QIAamp RNA blood minikit (Qiagen, Valencia, CA, USA). We performed rectal swabs using the QIAamp DNA stool minikit as well as the QIAamp Viral RNA minikit (Qiagen, Valencia, CA, USA) to extract DNA and RNA, respectively.

The manufacturer’s recommended procedures were followed for nucleic acid extraction. Reverse transcription of total RNA to cDNA was performed with the iScript cDNA synthesis kit (Bio-Rad, Hercules, CA) following the manufacturer’s instructions.

We selected a consensus primer for each viral pathogen to avoid possible genetic divergence of pathogens in wildlife, which cannot be amplified by a specific primer designed for analyzing domestic animals (21). Samples and primers selected for (RT)PCR screening are listed in Table 1. The limitation of detection of (RT)PCR under designed conditions for amplifying the genes of targeted infectious agents ranged from 1 to 1000 gene copies/µL (Table 2).

**TABLE 2.** Sensitivity of specific PCR assays for detecting CPPV-1, FeLV, FIV, CoV, and CDV. The target genes were cloned into a plasmid vector and the plasmid was diluted to 10^0^ to 10^9^ gene copies/µL for each detection assay

| Targeted<br>agent | Sensitivity (Gene copies/μl) |
| --- | --- |
| CPPV-1 | 10 gene copies/μl |
| FeLV | 100 gene copies/μl |
| FIV | 10 gene copies/μl |
| CoV | 100 gene copies/μl |
| CDV | 10 gene copies/μl |

The PCR amplicons of collected samples were sequenced in an ABI377 sequencer using an ABI PRISM dye-terminator cycle sequencing ready reaction kit with Amplitaq DNA polymerase (Perkin-Elmer, Applied Biosystems). To identify sequences similar to those of the amplicons, a BLAST search was performed using GeneBank with the nt/nr database available on the BLAST website (BLAST; https://blast.ncbi.nlm.nih.gov/Blast.cgi).

### Phylogenetic analysis

The nucleotide sequences of the infectious agents amplified in this study and retrieved from NCBI Genbank (https://www.ncbi.nlm.nih.gov/nucleotide/) accorded with CLUSTALW (28) in the MEGA 7 software program (29). The maximum-likelihood method (30) was used to model the phylogenetic relationship among sequences amplified from each infectious agent. Prior to the construction of a maximum-likelihood tree, the most suitable model was determined using MEGA 7 based on the lowest Bayesian information criterion (BIC) score (31).

### Data analysis

We first estimated the prevalence of each targeted infectious agent and its 95% confidence interval (CI) (32). As leopard cats are endangered, our sample size was limited; thus, we did not intend to exclude the possible distribution of the targeted infectious agents in the population of leopard cats if all individual samples screened negative.

The samples from live-trapped (LT) and FD leopard cats were pooled to evaluate a possible spatial or temporal cluster of target pathogens using SaTScan version 9 (33) with the Bernoulli model (34).

## RESULTS

### Leopard cat sample collection and distribution in Miaoli County

From 2015 to 2019, we collected samples from 52 leopard cats, of which 23 were LT and 29 were FD (Table 3; Table S1). No significant difference in sex was noted between LT and FD individuals (Pearson’s chi-squared test; p = 0.157). However, there were significantly more adults in the FD group than in the LT group (Fisher’s exact test; p = 0.0026). Samples were collected from leopard cats across western Miaoli County in a landscape of fragmented secondary forest habitat surrounded by farmland and residential areas (Fig. 1), which corresponded to the current distribution of the leopard cat population.

**TABLE 3.** Sex and age classification of leopard cats collected from live-trapped and found-dead individuals

| **Type of animal**<br>**analyzed** | **Female (n = 19)** |  |  | **Male (n = 33)** |  |  | **Total** |
| --- | --- | --- | --- | --- | --- | --- | --- |
|  | **Adult** | **Subadult** | **Juvenile** | **Adult** | **Subadult** | **Juvenile** |  |
| Live-trapped | 3 | 4 | 4 | 4 | 8 | 0 | 23 |
| Road killed | 6 | 2 | 0 | 16 | 4 | 1 | 29 |
| Total | 9 | 6 | 4 | 20 | 12 | 1 | 52 |

### Prevalence and distribution of targeted viral pathogens

For the targeted viral pathogens, only CPPV-1 and coronavirus were detected in the collected samples of leopard cats. The prevalences (95% CI) of CPPV-1, FeLV, FIV, CoV, and CMV were 63.5% (50.4%–76.6%), 0% (0%–6%), 0% (0%–5.9%), 8.8% (0%–18.4%), and 0% (0%–6.3%), respectively (Table 4). The prevalence of CPPV-1 in FD cats was significantly higher than that in LT cats (Fisher’s exact test, p = 0.002). Furthermore, the prevalence was significantly higher in adults than in subadults (Fisher’s exact test, p = 0.01). We did not determine any difference in prevalence between the type of sample, sex, and age for CoV (Table 4).

**TABLE 4.**
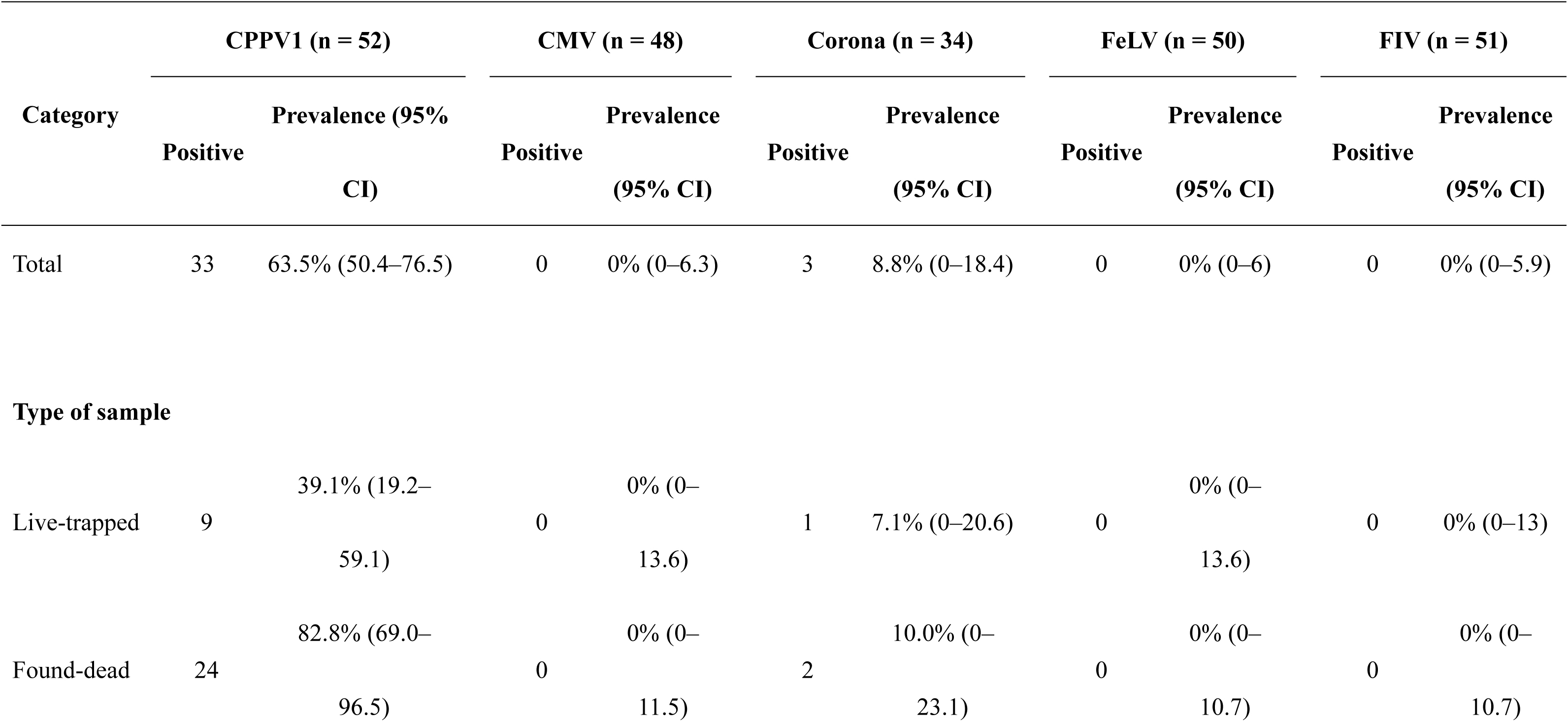
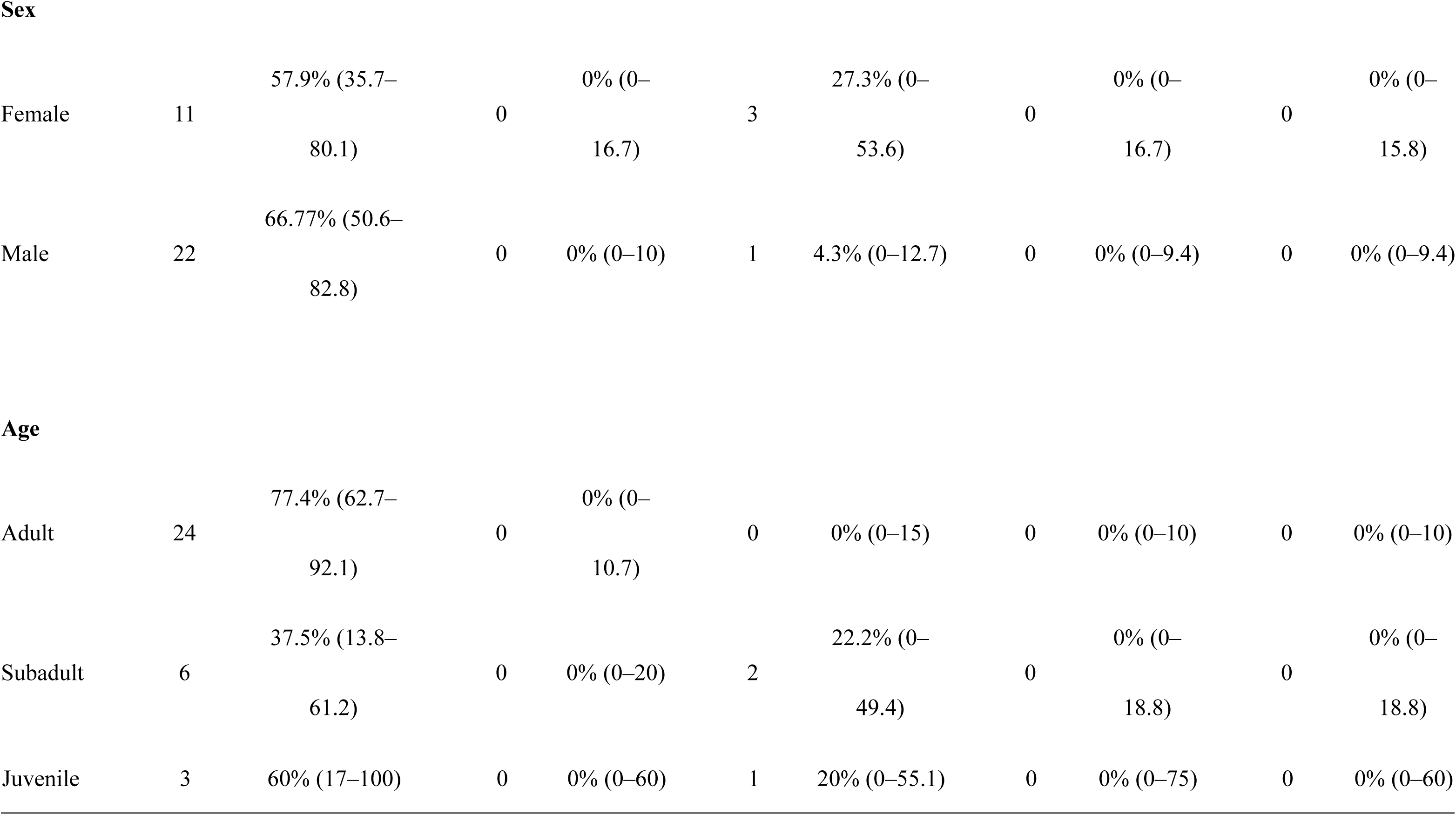
Prevalence of targeted viral pathogens in the free-living leopard cat population according to sample type, sex, and age

| Category | CPPV1 (n = 52) |  | CMV (n = 48) |  | Corona (n = 34) |  | FeLV (n = 50) |  | FIV (n = 51) |  |
| --- | --- | --- | --- | --- | --- | --- | --- | --- | --- | --- |
|  | Positive | Prevalence (95% CI) | Positive | Prevalence (95% CI) | Positive | Prevalence (95% CI) | Positive | Prevalence (95% CI) | Positive | Prevalence (95% CI) |
| Total | 33 | 63.5% (50.4–76.5) | 0 | 0% (0–6.3) | 3 | 8.8% (0–18.4) | 0 | 0% (0–6) | 0 | 0% (0–5.9) |
| Type of sample |  |  |  |  |  |  |  |  |  |  |
| Live-trapped | 9 | 39.1% (19.2–59.1) | 0 | 0% (0–13.6) | 1 | 7.1% (0–20.6) | 0 | 0% (0–13.6) | 0 | 0% (0–13) |
| Found-dead | 24 | 82.8% (69.0–96.5) | 0 | 0% (0–11.5) | 2 | 10.0% (0–23.1) | 0 | 0% (0–10.7) | 0 | 0% (0–10.7) |
| Sex |  |  |  |  |  |  |  |  |  |  |
| Female | 11 | 57.9% (35.7–80.1) | 0 | 0% (0–16.7) | 3 | 27.3% (0–53.6) | 0 | 0% (0–16.7) | 0 | 0% (0–15.8) |
| Male | 22 | 66.77% (50.6–82.8) | 0 | 0% (0–10) | 1 | 4.3% (0–12.7) | 0 | 0% (0–9.4) | 0 | 0% (0–9.4) |
| Age |  |  |  |  |  |  |  |  |  |  |
| Adult | 24 | 77.4% (62.7–92.1) | 0 | 0% (0–10.7) | 0 | 0% (0–15) | 0 | 0% (0–10) | 0 | 0% (0–10) |
| Subadult | 6 | 37.5% (13.8–61.2) | 0 | 0% (0–20) | 2 | 22.2% (0–49.4) | 0 | 0% (0–18.8) | 0 | 0% (0–18.8) |
| Juvenile | 3 | 60% (17–100) | 0 | 0% (0–60) | 1 | 20% (0–55.1) | 0 | 0% (0–75) | 0 | 0% (0–60) |

The spatial distribution of CPPV-1-positive individuals was scattered in the west of Miaoli County. Three positive CoV samples were distributed in northwest Miaoli (Fig. 2). We did not determine any spatial and temporal aggregation of CPPV-1 infection in the sampling area (SaTScan, Bernoulli model, p = 0.094). Spatial and temporal analyses were not performed for CoV, CMV, FeLV, and FIV, because very few or no positive samples were detected.

**FIG 2.**
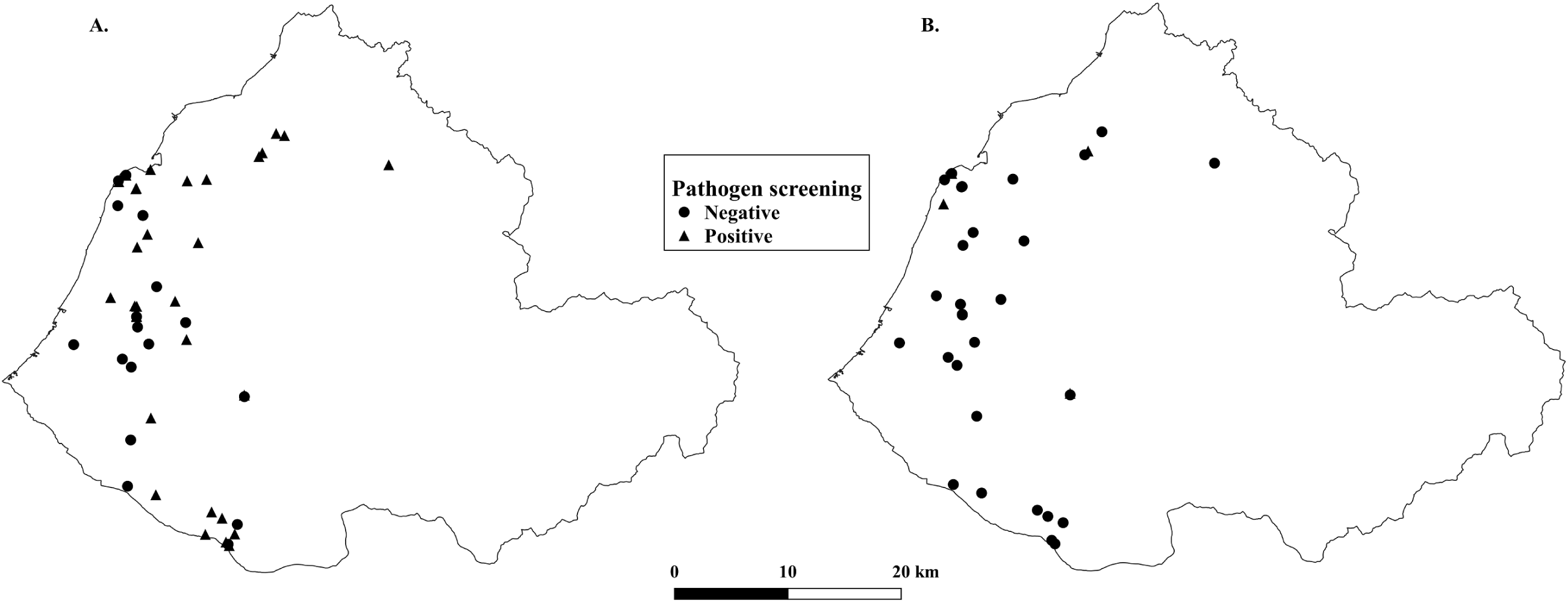
Spatial distribution of CPPV-1(A) and CoV (B) in leopard cats. No significant aggregation of positive samples was noted for either CPPV-1 or CoV.

### Viral strain identification and phylogenetic analysis

Viral strain identification of CPPV-1 was based on the VP2 amino acid sequences obtained from the 29 CPPV-1-positive leopard cats. We determined that 11, 7, 6, and 5 leopard cats were infected with CPV-2a, CPV-2b, CPV-2c, and feline panleukopenia virus (FPV), respectively (Table S2). The occurrence of CPPV-1 strain was significantly different from 2015 to 2018 (Fisher’s exact test, p = 0.006), with CPV-2b occurrence decreasing and CPV-2c and FPV increasing (Fig. 3).

**FIG 3.**
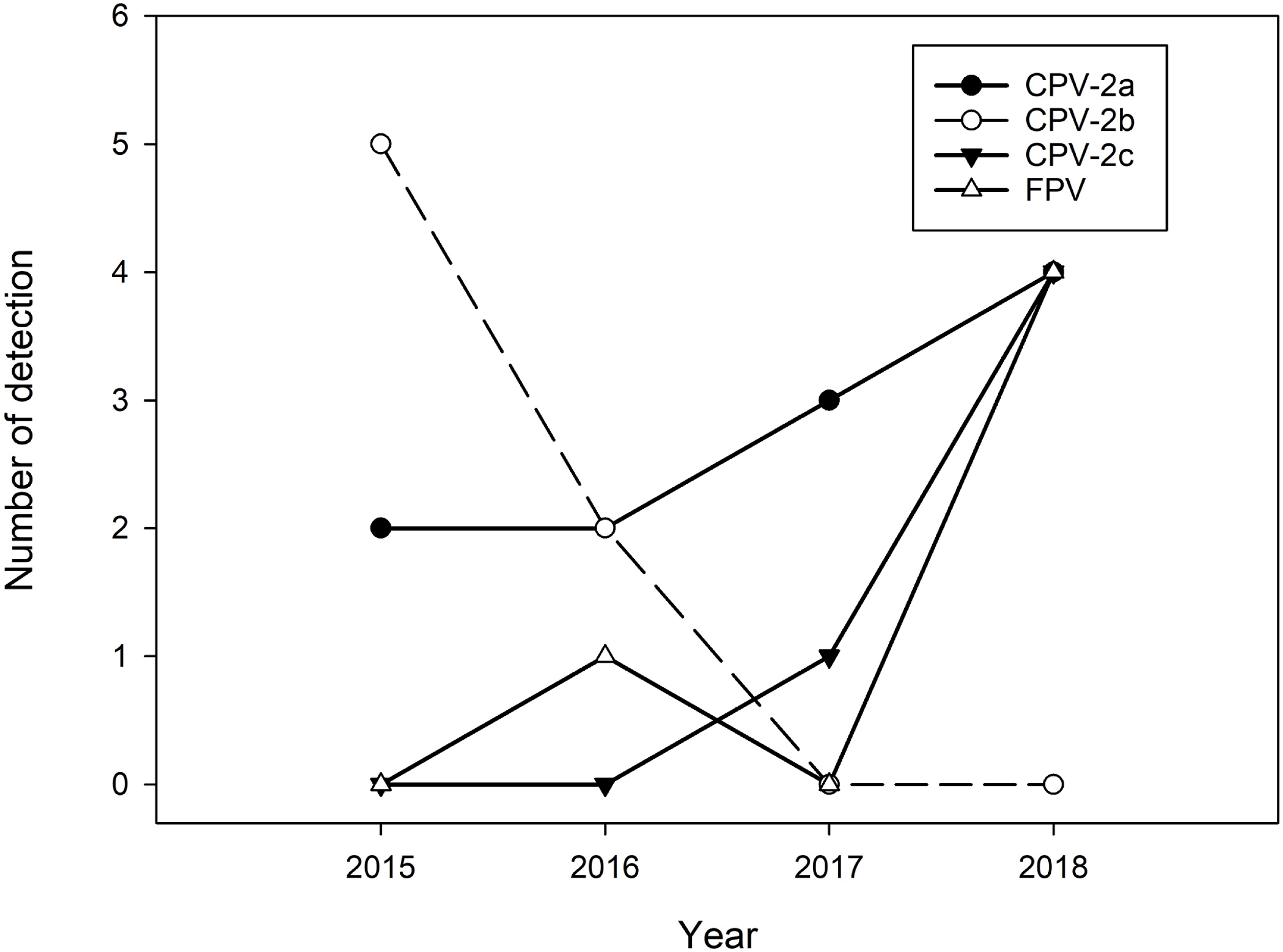
Frequency of positive detection for CPPV-1 strains from 2015 to 2018. The detection of CPV-2b decreased with an increase in CPV-2c and FPV detection.

Partial VP2 sequences of all CPPV-1 strains amplified from 29 leopard cats, 27 dogs, and 9 cats in Miaoli County and accessed from Genbank were included for phylogenetic analysis (Table S3). We adopted the Tamura-Nei model to construct a CPPV-1 phylogenetic tree based on the lowest BIC scores. The phylogenetic tree indicated that each CPPV-1 strain amplified from leopard cats and domestic carnivores from Miaoli County was primarily located in the same subcluster (Fig. 4). Furthermore, the majority of sequences of each CPPV-1 strain amplified from Taiwanese leopard cats and domestic carnivores were identical, comprising sequence types CPV-2a/1, CPV-2b/5, CPV-2b/8, CPV-2c/3, and FPV-4 (Fig 4, Table S3). However, certain sequence types were detected in leopard cats but not in domestic carnivores (Fig. 4, Table S3). Most of the nucleotide mutations of different CPPV-1 variants amplified from leopard cats were synonyms, which did not change the encoded amino acid (Fig 4; Table S2). Nonsynonymous mutations of sequence types amplified from leopard cats were determined in CPV-2a/3 with P352L and P356S substitution, CPV-2b/7 with S339N substitution, CPV-2c/5 with G437E substitution, FPV/5 with A379V substitution, and FPV/6 with Q310L, A334T, R377K, or R382K substitution.

**FIG 4.**
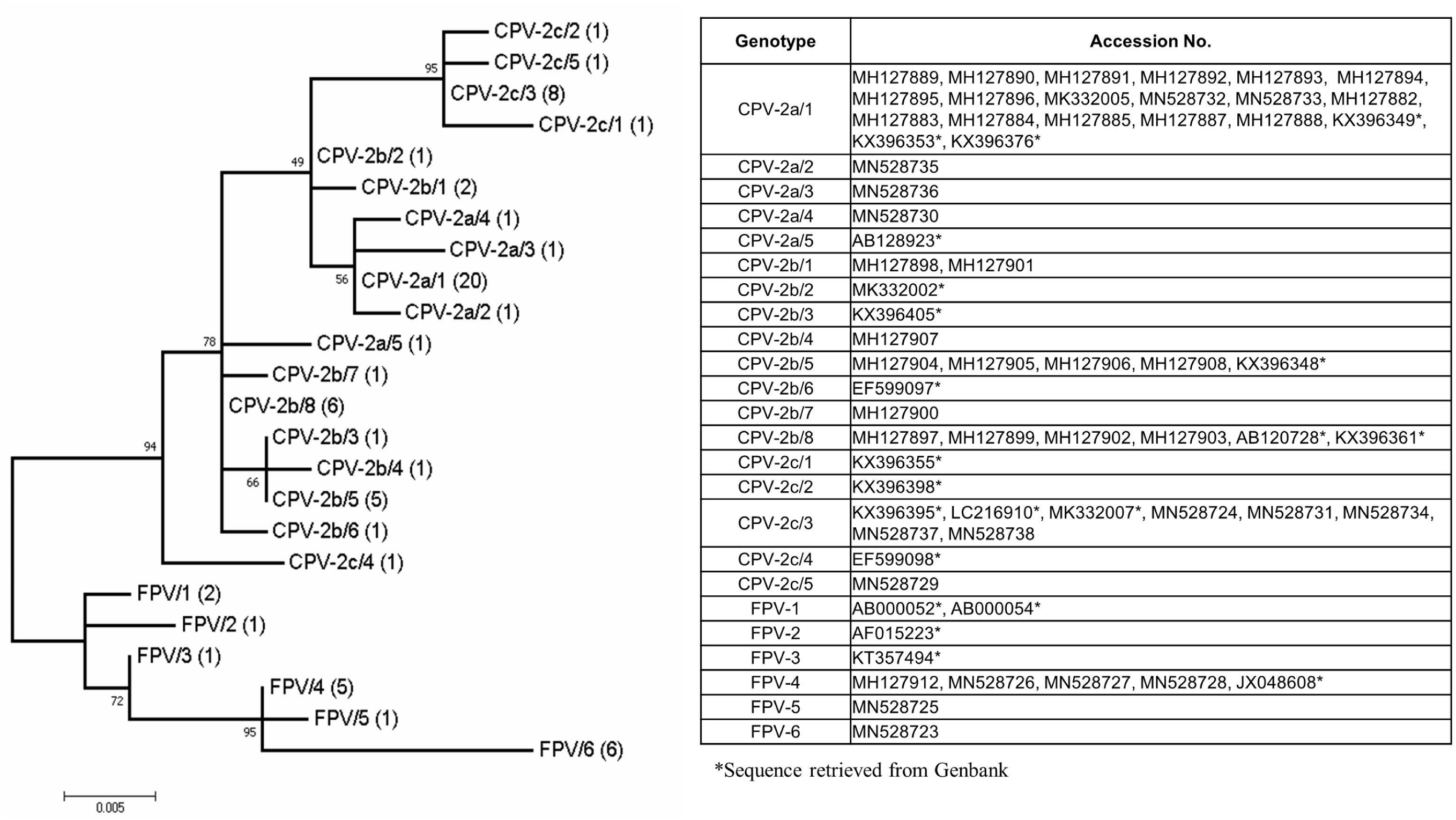
Molecular phylogenetic relationship of the partial VP2 sequences of the Carnivore protoparvovirus 1 amplified from leopard cats, domestic carnivores, and sequences retrieved from GenBank. The bootstrap value is reported next to the node with 1,000 replicates. Each strain and sequence type is labeled and followed by the number of identical sequences within each group (e.g., CPV-2a/1 (19), indicating that the sequence type 1 of the CPV-2a strain contains 19 identical sequences). The host species and location of the isolates of each accession number was assessed (Table S3).

Phylogenetic analysis of the 3 sequences amplified from the RNA-dependent DNA polymerase (RdRP) gene of CoV from leopard cats was first performed using the Tamura 3-parameter model with discrete Gamma distribution. The phylogenetic tree indicated that all the amplified CoV sequences from leopard cats were located in a cluster of viral species, Alphacoronavirus 1, and a feline coronavirus subcluster (Fig. 5).

**FIG 5.**
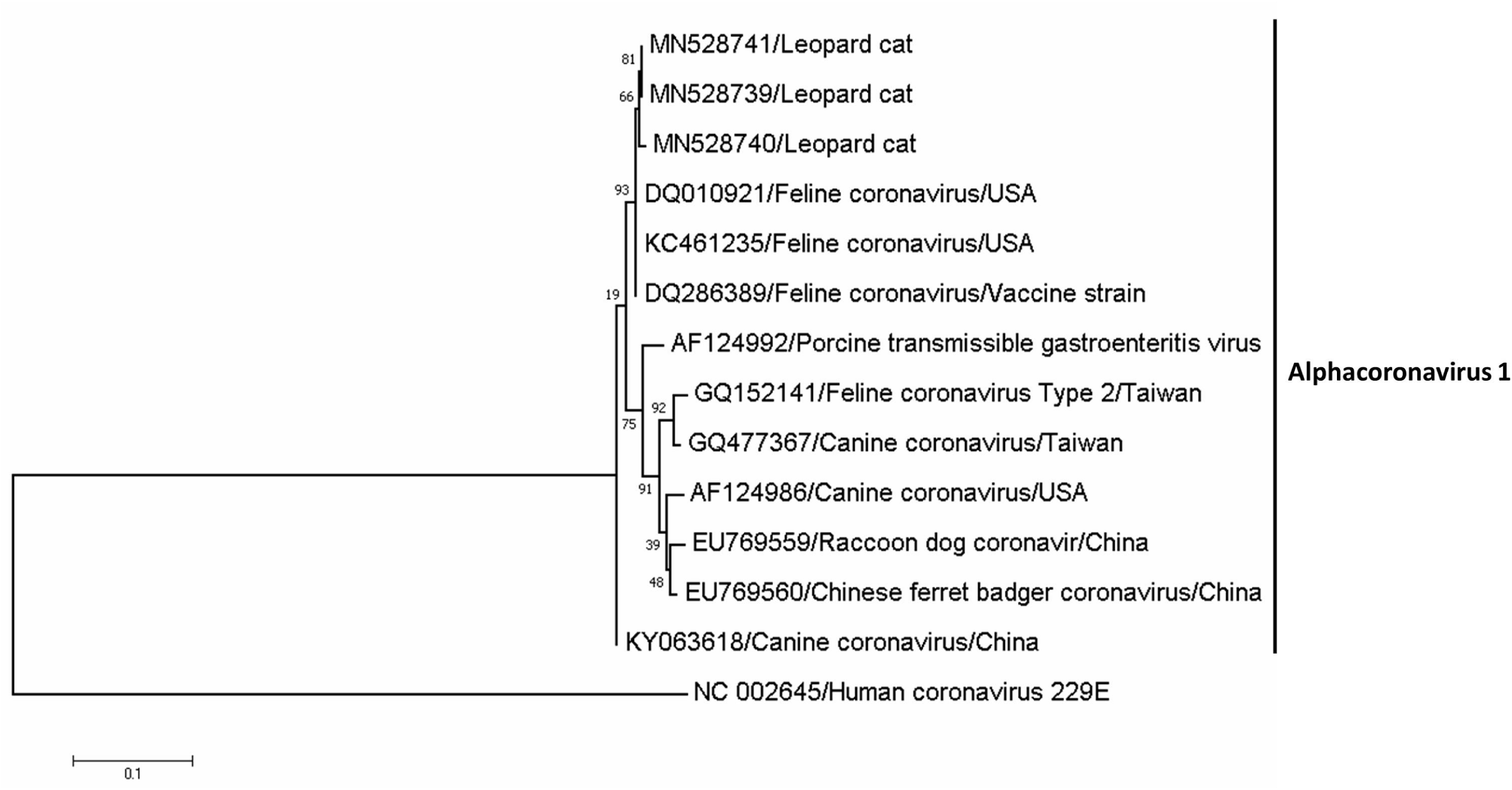
Molecular phylogenetic relationship of the partial RNA-dependent RNA polymerase gene of coronavirus amplified from leopard cats, and sequences retrieved from GenBank. The bootstrap value is reported next to the node with 1,000 replicates. Three amplified sequences for leopard cats (Genbank accession number: MN528739 – MN528741) were located in the feline coronavirus cluster.

## DISCUSSION

In this study, we screened the selected viral pathogens using (RT)PCR and determined the distribution of CPPV-1 and CoV in free-living leopard cats. Phylogenetic analysis revealed that the majority of identical genetic types of CPPV-1 strains were circulated between leopard cats and domestic carnivores; however, unique genetic types were identified in leopard cats. On the basis of the sequences of the RdRp gene, all the amplified CoV strains were identified as strains of feline coronavirus (FCoV) in species of Alphacoronavirus 1.

To our knowledge, CPPV-1 and FCoV infection in free-living leopard cats has only been reported in Taiwan (6), although CPPV-1 infection has been previously reported in captive leopard cats from Taiwan and Vietnam (11, 12). The worldwide distribution of CPPV-1 has resulted in the infection of various wild carnivorous species (22, 35–38). Mech et al. (38) determined that CPPV-1 contributed to a 40% to 60% reduction in wolf pup survival and impeded the population growth rate. Disease induced by CPPV-1 infection was commonly found in the juvenile or subadult individuals of domestic carnivores. However, adult individuals with severe clinical signs of CPPV-1 infection were recorded (39–41). Studies are increasingly reporting severe CPPV-1 enteritis in adult dogs (40, 42). Furthermore, a higher risk of developing chronic gastrointestinal disease had been determined in dogs after CPPV-1 infection (43). However, we observed a higher prevalence of CPPV-1 in FD and adults. A higher prevalence may represent a higher risk of infection or lower mortality. Prevalence data alone are not sufficient to evaluate the effect of CPPV-1 on different sample types or age categories. Therefore information regarding the physical effects, pathological changes, and mortality caused by CPPV-1 is required.

FCoV infection has been documented in various domestic and wild felids (44–47). The infection can be asymptomatic or associated with a fatal systematic disease, feline infectious peritonitis (FIP), and enteric disease (48, 49). Mochizuki et al. (50) screened serum antibodies of 17 iriomote cats (*Prionailurus bengalensis iriomotensis*), a subspecies of leopard cats, for coronavirus and found a prevalence of 82%. This study indicated frequent exposure to and transmission of FCoV in leopard cats. Although FCoV is commonly detected in wild felids worldwide, only a few species, such as cheetahs (*Acinoyx jubatus*), have been reported to exhibit FIP (44, 46, 48). In our study, 2 out of 3 positive samples were from FD cats and 1 was from an LT cat. We did not determine any pathological changes or clinical signs related to FCoV. Nevertheless, felids infected with FCoV that display no evidence of disease are considered to be chronic carriers that may increase other felids’ risk of contracting FIP (45, 49).

In this study, the effects of CPPV-1 and FCoV on individuals or the population of these leopard cats were not evaluated. However, based on the documented effects and cases of CPPV-1 and FCoV on wild felids, the effect of CPPV-1 and FCoV on leopard cats should not be overlooked, and continuous surveillance will be required.

Moreover, the spatial aggregation of CPPV-1 infection in leopard cats was not determined in the sampling area, which indicated a wide distribution of CPPV-1 in the habitat of leopard cats. CPPV-1 is stable in the environment and infectiousness can be maintained for several months (35). Free-roaming domestic carnivores are commonly observed in the sampling area, which is an active area with well-developed road systems (51, 52). Although the sample size was small, we found a very high prevalence of CPPV-1 (90%; n = 10; data not shown) in free-roaming dogs and cats in our sampling area. These conditions aggravate the transmission and distribution of CPPV-1 in the leopard cat habitat. Future studies should evaluate the influence of domestic carnivores on the transmission of CPPV-1 in the habitat.

In addition to pathogen surveillance, application of molecular analysis techniques for pathogens has been suggested for investigating several aspects of pathogenesis (53), including pathogen characterization and pathogen transmission (53). We identified the infection of 4 strains of CPPV-1 and FCoV in leopard cats based on the sequences of each positive amplification for selected pathogens. Temporal dynamics revealed that the infection of CPV-2c and FPV was increased, whereas CPV-2b infection was decreased. The distribution of CPV-2c in Taiwan was first detected in dogs in 2015 (54). Since then, CPV-2c has gradually become the predominant variant of CPPV-1 in dogs (54). We first detected CPV-2c in leopard cats in 2017, which indicates an original transmission direction of CPPV-1 from domestic carnivores to leopard cats. Background information and surveillance data for FPV are scarce. Therefore, factors that increase FPV infection rates still need to be assessed.

Our previous study found that the majority of sequences of CPPV-1 variants were identical between domestic carnivores and the leopard cats based on the partial VP2 gene sequences (6), which suggested frequent transmission of CPPV-1 between domestic and wild carnivores. In this study, we collected 2 times of leopard cat samples and we recorded several different sequence types for each CPPV-1 variant circulating in the leopard cat population (Fig. 4). However, the majority of amplifications from both domestic carnivores and leopard cats belonged to a specific sequence type of each variant. These results support the assumption that CPPV-1 is transmitted between domestic carnivores and leopard cats. Although we identified nonsynonymous mutations of sequence types from leopard cats, the causes and function of amino acid substitutions were undetermined. The amplified DNA sequence of CPPV-1 VP2 encoded amino acid from 300 to 437 residues (Table S2), located in the GH loop, an externally exposed loop in the antigenic region with the greatest variability (55). The sequence types of each variant found only in leopard cats does not indicate an adoption to the leopard cat, as only a few sequences from domestic carnivores in the sampling area were reported.

Cross-species transmission of CPPV-1 between domestic and free-living carnivores has been demonstrated or suspected in several countries (35, 36). Due to the critically endangered situation of leopard cats in Taiwan, sustained CPPV-1 transmission in this low-density population is improbable (56). We considered the domestic carnivores as the primary reservoirs based on the evidence that dogs and cats exhibited the highest abundance among carnivores in the study area, a high prevalence of CPPV-1, and the fact that CPV-2c occurrence in domestic dogs was earlier than in leopard cats.

In this study, we did not detect any current infection of FIV, FeLV, and CDV in leopard cats. The 95% CI of prevalence for FIV, FeLV, and CDV was 0% to 5.9%, 0% to 6%, and 0% to 6.3%, respectively. On the basis of the long-lasting disease and proviral DNA in peripheral blood monocytic cells characteristics of the FIV and FeLV in Felidae, the possibility of a false negative is low. Therefore we considered a low occurrence of FIV and FeLV in Taiwanese leopard cats. Studies have been conducted involving serological surveys of FIV and FeLV in leopard cats in Taiwan and Vietnam and did not detect any positive cases (12, 57). However, Hayama et al. (10) determined a prevalence of 3% (n = 86) and 13.6% (n = 280) of FIV infection in Tsushima leopard cats and domestic cats, respectively, in Kami-Shima, Tsushima Island, Japan. The domestic cat was considered as the reservoir of FIV in the Tsushima case (10, 58). The prevalence of FIV and FeLV varies between different felids and geographic regions (59).

The infection of CDV has been reported in both wild and domestic felids (59, 60). Ikeda et al. (61) reported a captive leopard cat in Taiwan having antibodies against CDV. Furthermore, a serological survey for CDV found a prevalence of 77.8% in wild Taiwanese leopard cats (62). Exposure to CDV in Taiwanese leopard cats is considered to be high. However, none of the leopard cats manifested clinical signs of CDV. Although we targeted amplifying the nucleotide sequence of CDV and identifying the strain from leopard cats, the low detection probability was expected because of a short virus shedding period. Furthermore, a diseased individual may reduce their activity and thus the probability that they would be sampled.

Our study revealed CPPV-1 and FCoV infection in free-living leopard cats. The sympatric domestic carnivores are considered the primary reservoir for the pathogens identified in our study. Although the effects of CPPV-1 and FCoV on individual leopard cats and populations of leopard cats were not evaluated in this study, we strongly recommend the establishment of programs to manage CPPV-1 and FCoV in the leopard cat habitat with an emphasis on vaccination programs and population control measures for free-roaming dogs and cats. Previous studies have indicated that because of antigenic differences among CPPV-1 variants, new vaccines that also provide protection against the CPV-2c variant may need to be developed (40, 63).

## ACKNOWLEDGMENTS

This study was supported by a grant from the Ministry of Science and Technology (MOST)(108-2313-B-020-001) to C.-C. Chen. We thank the field crew members, especially Dr. Esther van der Meer for her assistance in the sample collection. This manuscript was edited by Wallace Academic Editing. We declare no conflict of interest.

